# Gene regulatory mechanisms underlying the evolutionary loss of a polygenic trait

**DOI:** 10.1101/307389

**Authors:** Mark Lammers, Ken Kraaijeveld, Janine Mariën, Jacintha Ellers

## Abstract

Trait loss is a pervasive phenomenon in evolution, yet the underlying molecular causes have been identified in only a handful of cases. Most of these involve loss-of-function mutations in one or more trait-specific genes. Parasitoid insects are fatty acid auxotrophs: they lost the ability to convert dietary sugars into fatty acids, a trait that is ubiquitous among non-parasitoid animals and is enabled by a highly conserved set of genes. Earlier research suggests that lack of lipogenesis is caused by changes in gene regulation, rather than gene decay. We compared transcriptome-wide responses to sugar-feeding in the non-lipogenic parasitoid species *Nasonia vitripennis* and the lipogenic *Drosophila melanogaster*. Both species adjusted their metabolism within four hours after feeding, but there was little overlap at the gene level between the responses of the two species. Even at the pathway-level, there were sharp differences between the expression profiles of the two species, especially in carbohydrate and lipid metabolic pathways. Several genes coding for key enzymes in acetyl-CoA metabolism, such as *malonyl-CoA decarboxylase* (MCD) and *HMG-CoA synthase* differed in expression between the two species. Their combined action likely blocks lipogenesis in the parasitoid species. Network analysis indicates most genes involved in these pathways to be highly connected, which suggest they have pleiotropic effects and could explain the absence of gene degradation. Our results indicate that modification of expression levels of only a few non-connected genes, such as MCD, is sufficient to enable complete loss of lipogenesis in *N. vitripennis*.

## Introduction

In recent years, there has been a growing awareness of the adaptive role of trait loss in evolution (Fong et al. 1995; Lahti et al. 2009; Ellers et al. 2012; Royer et al. 2016). Trait loss has been shown to affect resource use efficiency (D’Souza et al. 2014), speciation (Stoks et al. 2003), host-parasitoid co-evolution (Pascoal et al. 2016) and the evolutionary potential of lineages (Bejder and Hall 2002; McLean et al. 2011).

The molecular changes underlying trait loss have been resolved in only a small number of cases. In a number of these, one or several key genes of the underlying pathway have degraded. For example, loss of vitamin C production in primates is caused by a single frameshift mutation in the gene *Gulo* (Chatterjee 1973; Ohta and Nishikimi 1999). Loss of four opsin genes, combined with reduced expression of nine important transcriptional factors, underlies eye vestigialization in cave fish (Yang et al. 2016). Accumulation of frameshift mutations in enamel-specific genes was found in a number of toothless and enamelless mammal lineages (Meredith et al. 2011; 2013). These studies reveal the molecular changes underlying trait loss for particular traits with relatively simple genetic bases. However, the genomic basis of trait loss in more complex, polygenic traits is still unresolved.

The expression of a complex trait is generally regulated by a network with many interconnected pathways, containing intermediate steps encoded by a large number of genes. Trait loss can be the result of a disruption in any of the intermediate steps, but gene network theory predicts that degradation of highly connected genes (so-called hub genes) is usually detrimental or even lethal (Jeong et al. 2001; Lee et al. 2008; Cho et al. 2012; Winterbach et al. 2013) due to pleiotropic functions of these genes. Other studies show that gene pleiotropy also correlates to evolutionary constraints on gene expression (Papakostas et al. 2014; Morandin et al. 2017; Yang and Wittkopp 2017). Therefore, these genes are unlikely targets for molecular changes underlying decay of complex traits. In contrast, genes with a more peripheral position in the gene network have lower connectivity and fewer pleiotropic functions (He and Zhang 2006; Zhu et al. 2007). When such genes decay or their expression is suppressed, relatively few pleiotropic effects would result on other functions (reviewed in Winterbach et al. 2013). Regulatory changes might be a common mechanism underlying trait loss, but the role of gene expression and pleiotropic function in relation to trait loss is poorly understood (Gompel and Prud’homme 2009).

*De novo* synthesis of fatty acids is a highly conserved complex process involving many deeply conserved and highly pleiotropic genes (Towle et al. 1997; Letunic et al. 2008; Arrese and Soulages 2010). It is an integral part of the life-history of most animals, enabling them to convert dietary carbohydrates to lipids, which allows storing energy for leaner times and for resource allocation to reproduction. Despite these essential functions of fatty acid synthesis, lack of lipogenesis has repeatedly evolved across the eukaryotic tree of life. For example, several lineages of fungi are fatty acid auxotrophs, caused by loss or degradation of the fatty acid synthase gene (Xu et al. 2007; Luginbuehl et al. 2017). Also, multiple insect lineages lack lipogenesis as shown by labelling studies (Giron and Casas 2003; Visser et al 2012; Visser et al 2017) or a lack of increase in adult fat reserves, despite feeding on sugar *ad libitum* (Ellers 1996; Visser and Ellers 2008). In insects, this recurrent loss of lipogenesis is phylogenetically linked to the parasitoid lifestyle: parasitoid clades of flies, beetles and wasps have lost lipogenesis independently (Visser et al. 2010). Phylogenetic analysis also shows that the ability to synthesize fatty acids has re-evolved independently in a number of parasitoid wasp species (Visser et al. 2010). The repeated regain of lipogenesis, as well as the central role of the lipogenic pathway in the carbon metabolism, suggests that the loss of lipogenesis is due to modification of gene expression rather than genetic changes in protein coding regions of the genome.

In this study, we aim to unravel the changes in gene expression underlying the loss of lipogenesis in the parasitoid wasp *Nasonia vitripennis*. In *N. vitripennis*, there is no apparent gene loss or pseudogenization in pathways related to carbohydrate and lipid: a screening of the genome did not reveal gene losses in pathways relevant to lipogenesis (Werren et al. 2010). Visser et al. (2012) showed that in *N. vitripennis* sugar feeding does not cause a transcriptional response in several key genes in fatty acid biosynthesis, whereas these are upregulated under the same conditions in *Drosophila melanogaster*. These findings suggest that lipogenesis is decoupled from other metabolic processes, sharply contrasting with regulation of these processes in other organisms (Towle et al. 1997; Zinke et al. 2002; Sassu et al. 2012).

We characterized transcriptomic changes in response to sugar-feeding in the non-lipogenic parasitoid *N. vitripennis*. No lipogenic strains of *N. vitripennis* are known, therefore we compared these changes to the response of a well-characterized representative lipogenic species, *D. melanogaster*. Both species have annotated genomes, well-studied life-histories and readily feed on sugar solutions. Although these species are in different insect orders, the metabolic pathways underlying lipid synthesis are highly conserved. We first compared the transcriptomic response to sugar-feeding for both species separately. Next, we compared these responses between the two species. Our results show that both species respond to sugar-feeding by adjusting transcription of key genes involved in multiple metabolic pathways, but network-based analyses indicate that both organisms evolved contrasting strategies in metabolizing dietary sugar.

## Material and Methods

### Study species

We used *Drosophila melanogaster* (Diptera: Drosophilidae) Bloomington stock 2057 (RRID:BDSC_2057, Adams 2000) and *Nasonia vitripennis* (Hymenoptera: Pteromalidae) strain AsymCX (Werren et al. 2010) in our experiments since the reference genomes originated from these strains. All strains were kept at 25°C, 75% relative humidity and 16:8h L:D cycle prior to and throughout the experiment.

### Experimental setup

All newly emerged insects were kept without access to food for 24-36h (only water). This pre-treatment period ensured that all were hungry and eager to feed. Next, mated females were randomly assigned to either of two treatments: starved (St) and sucrose-fed (Sc). Starved insects were kept without food for another 4h; sucrose-fed insects were given *ad libitum* access to a sucrose solution (20% w/v). All insects of the latter treatment were observed to feed within a few minutes. After exactly 4h all insects where killed by freezing in liquid nitrogen.

### Tissue collection and RNA extraction

Each experimental condition consisted of 10 individual females and the experiment was repeated three times, yielding three independent biological replicates. Frozen females were dissected on a clean liquid nitrogen-cooled steel block: only abdomens were retained for total RNA extraction. For *N. vitripennis*, eight females were pooled per RNA extraction. For *D. melanogaster*, each sample of RNA was extracted from two pools of four females which were subsequently combined. The remaining two individuals per replicate were stored frozen as backup. RNA was extracted using the Promega SV Total RNA Isolation System kit (Promega Corporation, USA) following manufacturer’s instructions except that RNA extracts were eluted in 30μL water. RNA concentrations were measured on a NanoDrop 2000 (Thermo Scientific) and the RNA Integrity Number was measured on a BioAnalyzer 2100 (Agilent). All samples had sufficient quantities of the required quality of RNA (supplementary table S1).

### Illumina sequencing

All samples were sent to the Beijing Genomics Institute where library preparation and sequencing were performed. This strand-specific TruSeq library preparation included poly-A RNA purification, mRNA fragmentation, cDNA synthesis from size-selected fragments using random primers and adapter ligation for sample identification. Resulting short-insert libraries were pooled and sequenced (90bp paired-end) on one lane of Illumina HiSeq2000. This yielded 13-20M reads per sample (detailed in supplementary table S1). Raw sequence data have been deposited in the NCBI Short Read Archive under the study accession number SRP127311.

### Transcriptomes and reference genomes

Sequence reads were checked for quality using FastQC version 0.10.1 [http://www.bioinformatics.babraham.ac.uk/projects/fastqc/]: only high quality clean reads were delivered. The reference genome annotations used for *N. vitripennis* and *D. melanogaster* were GCF000002325.3_Nvit_2.1 (downloaded from NCBI RefSeq 27 October 2014) and dmel_r6.11 (downloaded from FlyBase on 9 June 9 2016), respectively. Complete gene identifier conversion tables where created from the genome annotation files using a custom R script.

Indices to the reference genomes were built with Bowtie2-build version 2.2.4 [http://bowtie-bio.sf.net/bowtie2]. Reads were mapped to the respective reference genomes using TopHat2 version 2.0.13 (Kim et al. 2013) with the following settings: -r 0 -p 4 --library-type fr-firststrand. Statistics of mapping success are reported in supplementary table S1. Alignment files were converted to SAM-format using samtools version 0.1.19 (Li et al. 2009). GFF-files were converted to comply to HTSeq format requirements with a python script. Expression levels were counted per gene using HTSeq version 0.6.1 (Anders et al. 2015) with default settings. Only genes for which five or more libraries (for each species separately) had non-zero counts were retained because zero counts could cause spurious similarity between samples (Storey and Tibshirani 2003).

Differential gene expression analyses were performed in EdgeR version 3.10.2 (Robinson et al. 2010) as recommended by Guo et al. (2013). Gene expression was compared between the two treatments (starved vs. sucrose-fed) for each species separately. Normalization factors were calculated with the function calcNormFactors with default settings. A negative generalized log-linear model was fitted to the data and a Likelihood ratio test was used to obtain P-values (McCarthy et al. 2012). P-values were corrected for multiple testing using the R function p.adjust following the method of Benjamini-Hochberg (1995). A gene was considered significantly differentially expressed when the p-value associated with this comparison was below 0.05 after FDR correction.

### Gene Ontology Enrichment analyses

For each species, the set of differentially expressed genes was checked for enriched GO-terms with the R-package ‘topGO’ version 2.20.0 (Alexa et al. 2006). We used the weighed P-values that TopGO calculates using an algorithm that weighs statistical significances of higher GO categories by the significance of its lower categories: Higher categories are only recovered as significant when more genes are found at that category than expected by chance. GO-indexes for the *D. melanogaster* gene annotation were downloaded from FlyBase (Attrill et al. 2016). For *N. vitripennis* we used the Blast2GO-based GO-index for the Nvit2.1 assembly, available via DOI:10.13140/RG.2.1.2194.1929. Only GO-terms in the category ‘biological process’ were considered.

### Orthologous gene-based comparisons

Orthologs of *D. melanogaster* and *N. vitripennis* were obtained from OrthoDB7 (Waterhouse et al. 2013). This list of orthologous groups is based on previous versions of the *D. melanogaster* and *N. vitripennis* genome annotations and gene predictions that are not supported in the current annotations were omitted from the analysis. All single-copy orthologs were extracted from the database. Next, we calculated the Pearson’s correlation of the expression levels of the non-differentially expressed orthologs of *N. vitripennis* and *D. melanogaster*.

### Species-specific transcriptional responses

For each species, we manually checked the lists of differentially expressed genes for involvement in lipogenesis, regulation of lipogenesis, or carbohydrate metabolism pathways. Sets of differentially expressed genes between treatments were screened for regulatory components: (1) transcription factors, as listed in REGULATOR (Wang and Nishida 2015) and (2) non-coding RNAs, as annotated in the respective reference genomes.

Gene pleiotropy was estimated by the number of protein-protein interactions as collected in the database STRINGdb (Franceschini et al. 2013). The absolute change in expression level (logFC) in our treatments was tested for a correlation with the number of protein-protein interactions after log2 transformation.

### Visualization of KEGG-pathways

Each gene for which we obtained sufficient expression data was queried against the KEGG Pathway database (Kanehisa and Goto 2000; Kanehisa et al. 2008) in order to match it to a specific enzymatic reaction number. This analysis was repeated for *D. melanogaster* and *N. vitripennis* independently. These reactions were coupled to the corresponding gene expression data in our transcriptomes and imported into iPath2 (Letunic et al. 2008, http://pathways.embl.de/iPath2.cgi) to visualize the metabolic maps of both species. Our functional interpretation of biochemical pathways is based on Berg et al. (2006).

### Pathway-based comparison between transcriptomes of D. melanogaster and N. vitripennis

The number of DE genes per KEGG Pathway were summed for both species and divided by the total number of genes that each species had for that pathway. This fraction of induced genes per pathway is presented as a heatmap using the R function heatmap.2 from package gplots.

## Results

### Global gene expression patterns

We obtained 13.7-20.4 million high-quality 90bp paired-end reads for each of the 12 libraries (2 species, 2 treatments, 3 biological replicates). 17.7-24.2% of reads were unequivocally mapped to a single locus on the respective reference genomes and kept for subsequent analyses (supplementary table S1).

We first established that our treatment of four hours *ad libitum* access to sugar had a measurable effect on the insect’s abdominal transcriptome. Heatmaps of these transcriptomes are shown in figure 1. We found significant differences in expression level for 195 genes of *D. melanogaster* and 188 genes of *N. vitripennis* (table 1, supplementary tables S2 and S3). The number of enriched gene ontology terms (GO-terms) in the sets of DE genes are presented in table 1 and supplementary tables S4 and S5. 59 GO-terms were enriched in *D. melanogaster* and 19 in *N. vitripennis* (Fisher’s exact tests, weighted p-values < 0.05). These significantly enriched GO-terms contain a broad spectrum of processes altered upon sugar feeding, including terms related to amino acid metabolism, reproduction, and carbohydrate and fat metabolism.

**Figure 1.**
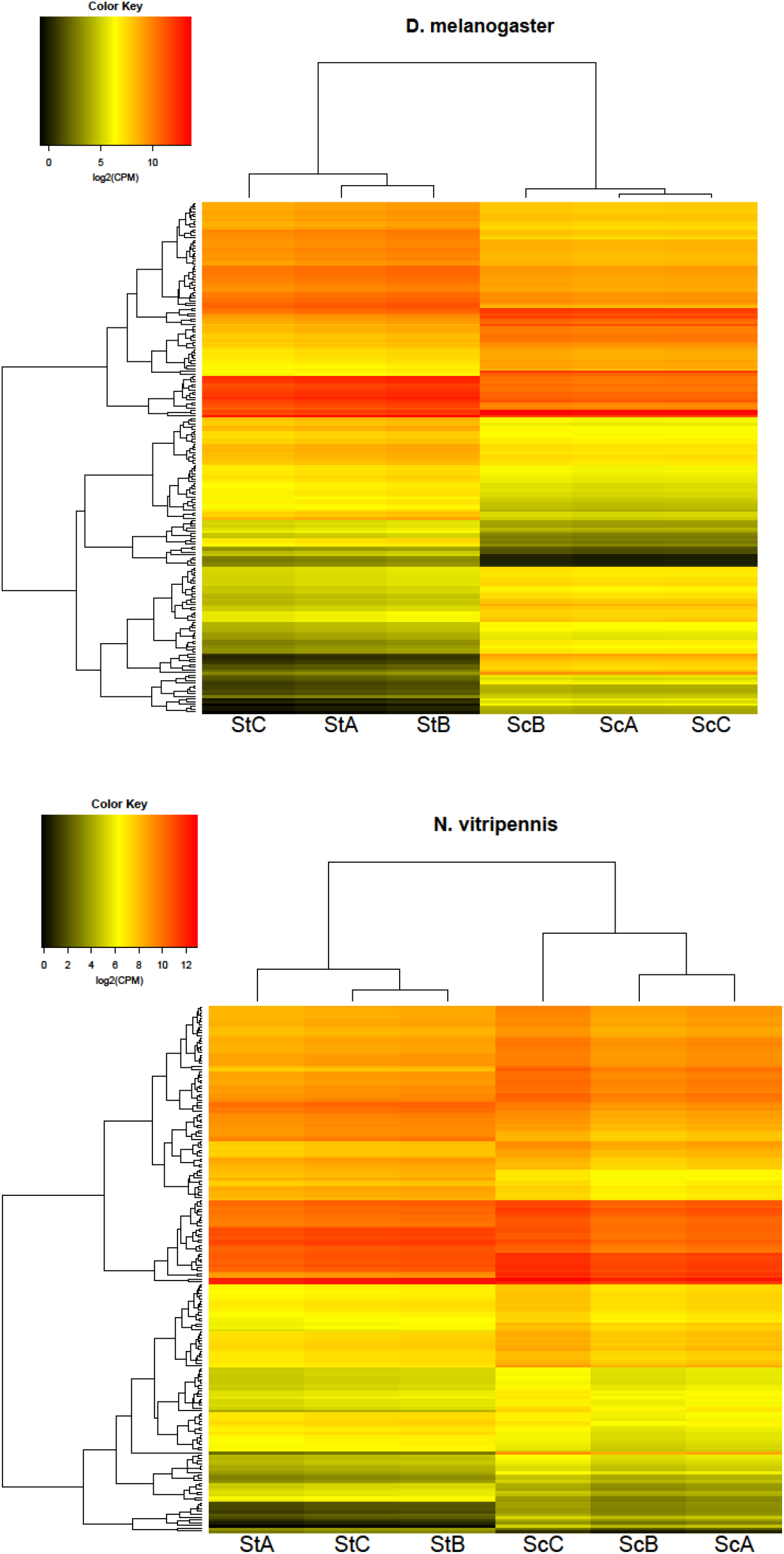
Heatmaps of the *D. melanogaster* and *N. vitripennis* transcriptomes. CPM = counts per million, St = starved, Sc = sucrose-fed, A/B/C = replicate.

**Table 1.** Summary of annotated genes for *D. melanogaster* and *N. vitripennis* and the number of genes up-and down-regulated upon sugar feeding with the number of unique Gene Ontology (GO) terms associated to these differentially expressed genes.

| Species | Number of genes in reference genome | Number of genes with expression data | Genes up-regulated | Genes down-regulated | GO terms |
| --- | --- | --- | --- | --- | --- |
| *D. melanogaster* | 17,974 | 12,977 (72.2%) | 84 | 111 | 59 |
| *N. vitripennis* | 14,321 | 10,914 (76.2%) | 126 | 62 | 19 |

To validate further comparison between the transcriptomic responses of these species, we compared the expression level of the 1822 non-differentially expressed single-copy orthologs of *D. melanogaster* against their expression level in *N. vitripennis* (figure 2). The overall gene expression patterns are positively correlated (Pearson’s r=0.58, t=30.155, df=1820, p<0.0001).

**Figure 2.**
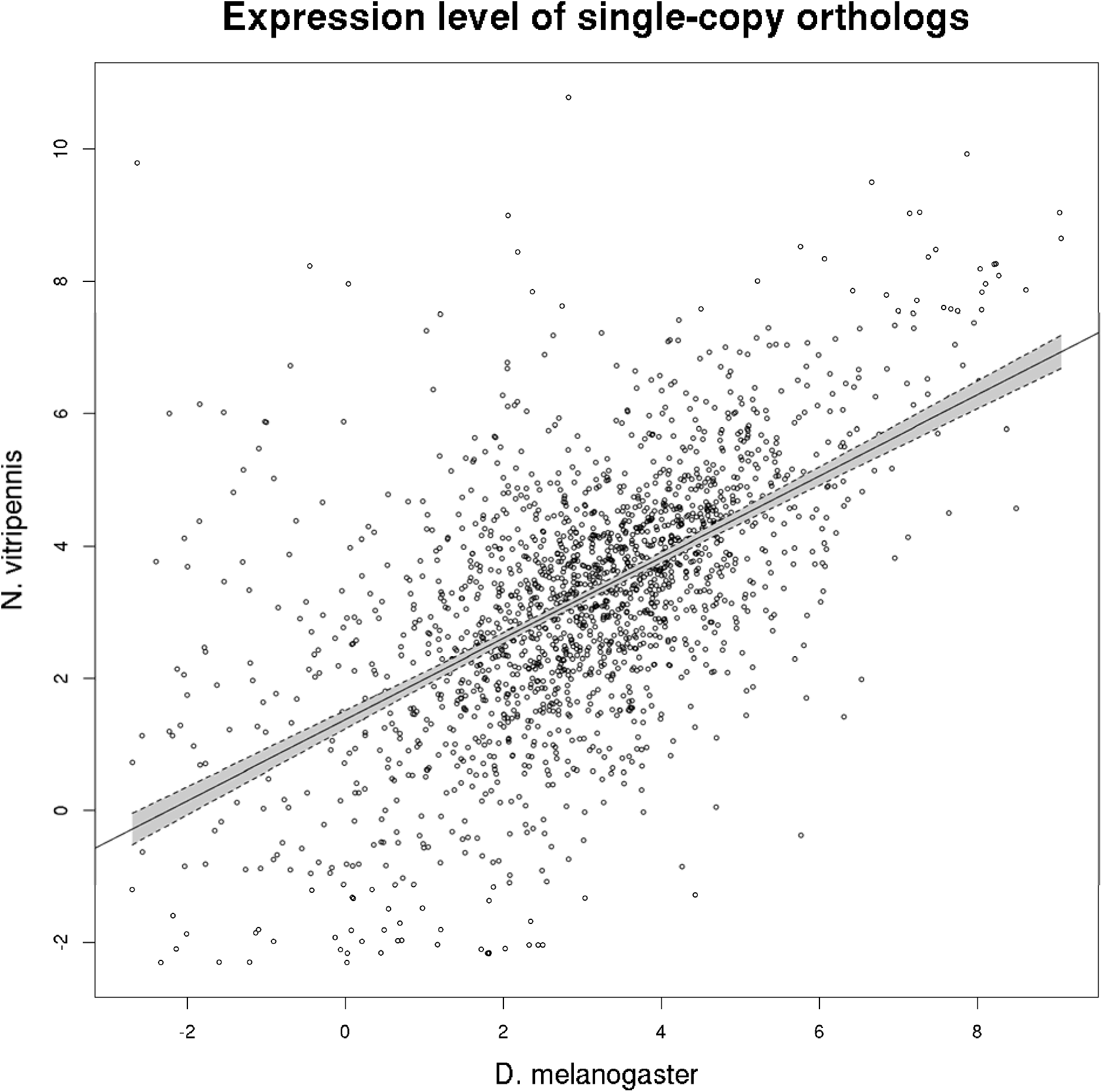
Interspecific correlation of expression level of non-differentially expressed single-copy orthologs between *D. melanogaster* and *N. vitripennis* when comparing sucrose feeding and brief starvation treatments.

### Transcriptional response to sugar-feeding in D. melanogaster

*D. melanogaster* upregulated *fatty acid synthase 1* (FBgn0283427) and *lipid storage droplet 1* (FBgn0039114) upon ingestion of sucrose. Genes related to catabolism of amino acids like *glutamate oxaloacetate transaminase 1* (FBgn0001124) were downregulated upon feeding as well. 1181 genes (11.0% of all genes for which we had expression data) were successfully matched with known reactions in KEGG Pathway. The resulting overview of the active and induced metabolic pathways is presented in supplementary figure S1. If we assume unaltered enzyme efficiencies and protein turnover rates overall, this figure provides a bird’s eye view on the organism’s abdominal metabolism. It shows that many processes other than our focal pathways were altered upon sugar-feeding, e.g. enzymes involved in nucleotide metabolism and amino acid metabolism.

Gene expression is regulated by a range of mechanisms, including non-coding RNAs and cis-regulatory components like transcription factors. Of the differently expressed genes, nine were annotated as non-coding RNAs (table 2). These were all down-regulated upon sugar-feeding. Only one of the differentially expressed genes was a transcription factor as listed in the transcription factor database REGULATOR: *sugarbabe*. It was strongly upregulated in our experiment.

**Table 2.**
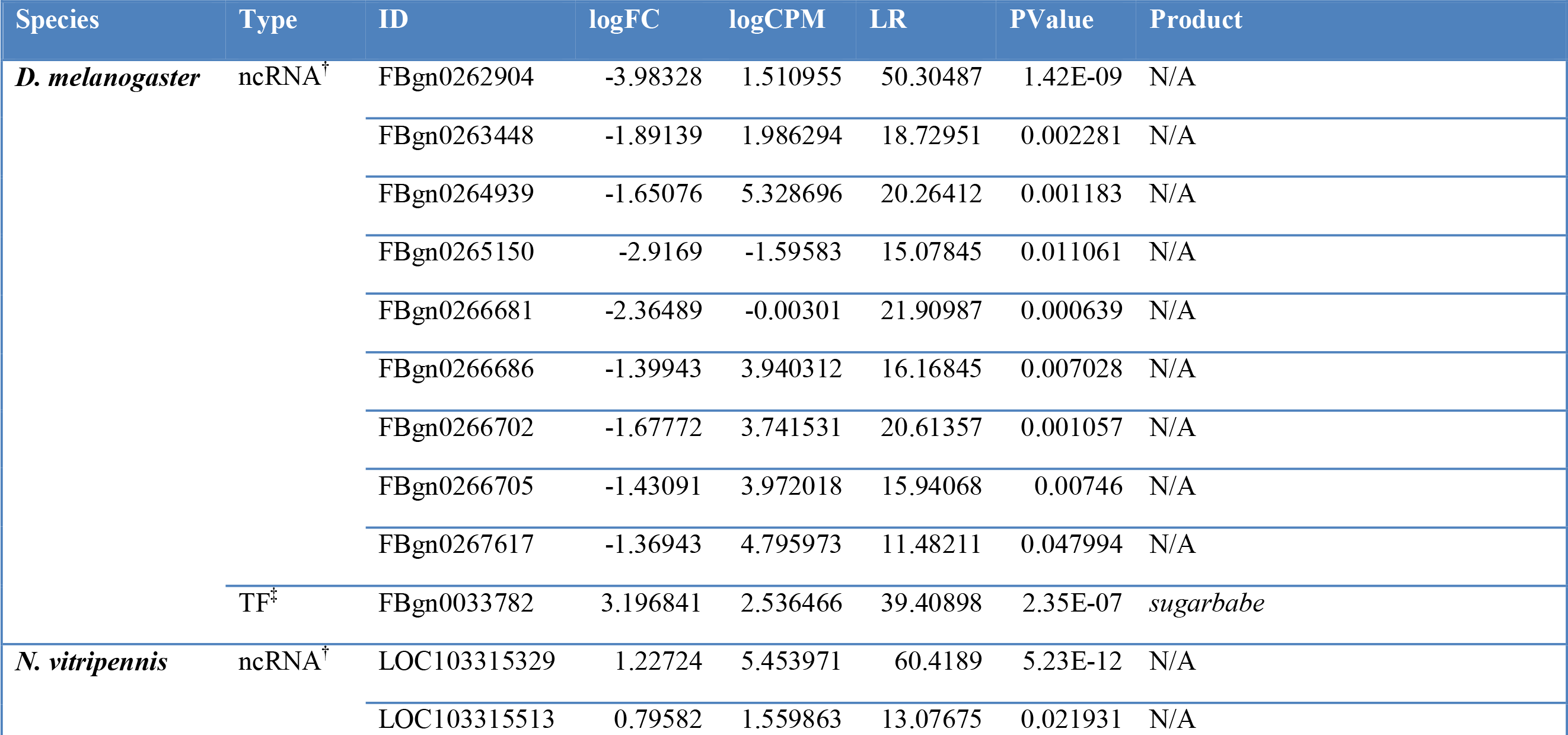
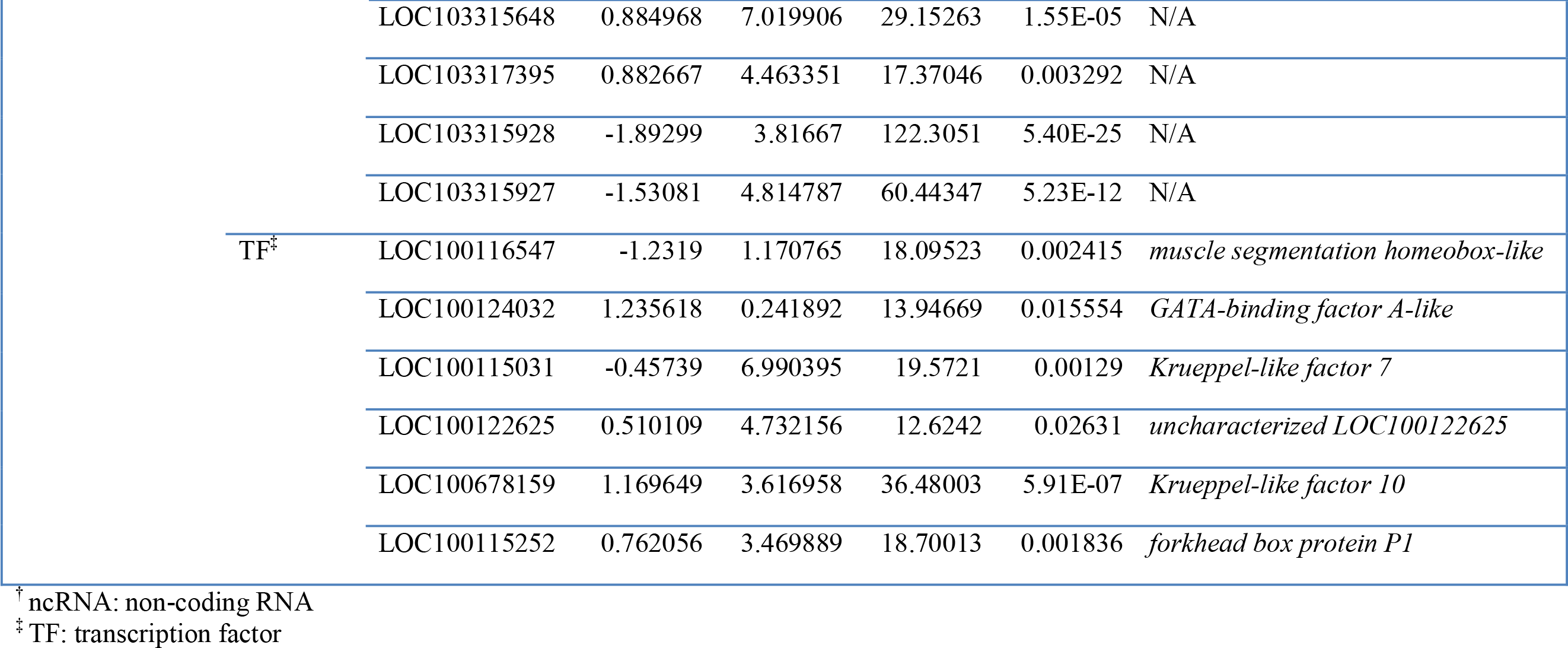
Differentially expressed regulatory genes for *D. melanogaster* and *N. vitripennis* in response to sugar feeding compared to a brief starvation treatment.

### Transcriptional response to sugar feeding in N. vitripennis

In *N. vitripennis* none of the three copies of the enzyme *fatty acid synthase* (LOC100121447, LOC100122099, LOC100122083) was upregulated in the sugar-fed treatment. Several other genes related to lipid metabolism were upregulated: *acetyl-CoA carboxylase* (LOC100123347), *glucose 6-phosphate dehydrogenase* (LOC100120232) and *ATP-citrate lyase* (LOC100119651) and a *citrate transporter* (LOC 100118210). A number of genes in other pathways linked to carbohydrate metabolism were differentially regulated upon sugar-feeding: *HMG-CoA synthase 1* (LOC100116401) and *phosphoenol pyruvate carboxykinase* (LOC10015526).

925 genes (8.5% of all genes for which we had expression data) had known reactions in KEGG Pathway. The resulting overview of the active and induced metabolic pathways is presented in supplementary figure S2, and the main changes of interest are summarized in figure 3.

**Figure 3.**
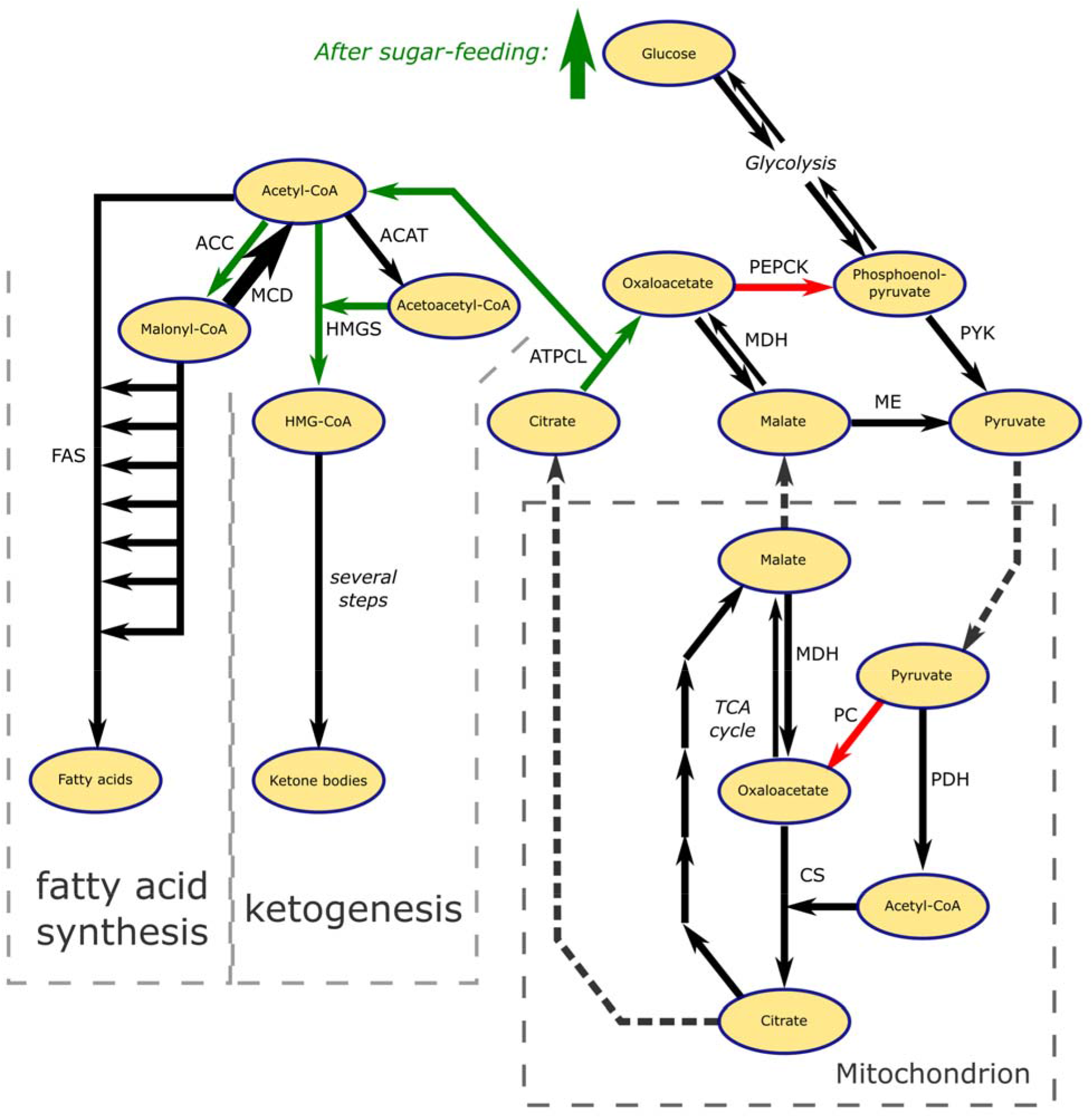
Differential gene expression in acetyl-CoA metabolism of *N. vitripennis* upon sugar feeding. Green and red arrows depict upregulated and downregulated genes, respectively. The thick arrow for MCD symbolize high constitutive expression. Dashed arrows represent transport of metabolites across the mitochondrial membrane.

Of the differently expressed genes in *N. vitripennis*, six genes were annotated as ncRNAs. Four of these were upregulated, two were downregulated (table 2). Of the transcription factors listed in REGULATOR, four were upregulated and two were downregulated expressed in our experiment (table 2).

### Comparison between N. vitripennis and D. melanogaster

While the majority of single-copy orthologs showed a strong correlation in expression level between *N. vitripennis* en *D. melanogaster* (see above), there were a number of outliers to this correlation: several genes were highly expressed in *N*. *vitripennis* at all times, but had a much lower expression in *D*. *melanogaster* (upper-left in figure 2). The genes with the most extreme interspecific difference in expression were *radish, Phosphodiesterase 1c, CG9117*, and *olf413. Radish* is a Rap-Like GTPase Activating Protein involved in memory dynamics (Folkers et al. 2006). *CG9117* is a metallo-beta-lactamase domain-containing protein without further annotation. *olf413* is a copper type II ascorbate-dependent monooxygenase. The opposite expression pattern (highly expressed in *D. melanogaster*, but low in *N. vitripennis)* was found for *Gasp* and *walrus. Gasp* is a chitin-binding protein associated with embryonic development. *walrus* is an electron transfer flavoprotein, probably capable of accepting electrons of several dehydrogenases. The full table of expression levels of non-plastic single-copy orthologs is available in supplementary table S6.

KEGG-pathways associated with the DE genes were visualised as a heatmap in figure 4 for both species. Full tables are available in supplementary table S7. In the carbohydrate metabolic pathways, *D. melanogaster* showed multiple DE genes in fructose (two genes), galactose (nine genes) and sucrose (eight genes) metabolism, while *N*. *vitripennis* had no genes differentially expressed in these pathways. By contrast, differentially expressed genes in *N*. *vitripennis* were involved in the tricarboxylic acid (TCA) cycle (four genes), propanoate and butyrate metabolism (two genes each) and pyruvate metabolism (three genes). There were also divergent responses in the amino acid metabolisms, lipid metabolic pathways and in the pathways of the metabolism of co-factors and vitamins.

**Figure 4.**
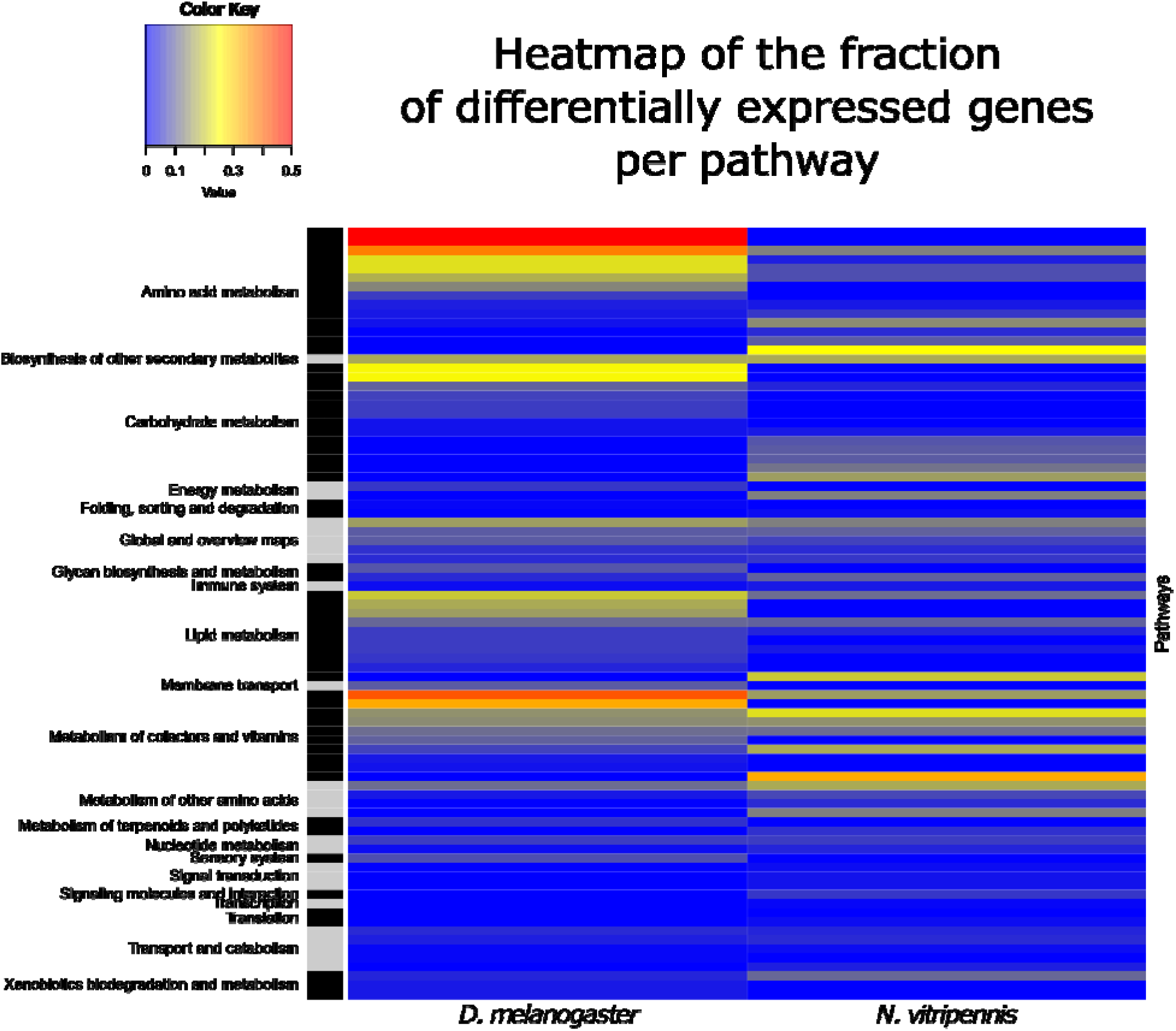
Heatmap of the fraction of genes per KEGG Pathway that was differentially expressed for *D. melanogaster* and *N. vitripennis* when comparing sucrose feeding and brief starvation treatments. The full table is provided in supplementary table S7.

### Differential expression correlates to gene pleiotropy

Figure 5 shows the correlation between absolute fold change of gene expression levels and the number of protein-protein interactions (PPI) as a measure for the level of pleiotropy. There was a significant negative correlation between fold change and the number of PPI: Pearson’s r = −0.195 for *N. vitripennis* (t=−20.772, df=10915, p<0.0001) and r = −0.216 for *D. melanogaster*(t=-25.192, df=12975, p<0.0001). Many genes related to fatty acid metabolism are positioned at the high end of the pleiotropy spectrum: the median number of PPI for genes from KEGG Pathway map ‘Fatty acid metabolism’ (map01212) is 16, while this number is only 11.5 for all metabolic pathways combined (map01100).

**Figure 5.**
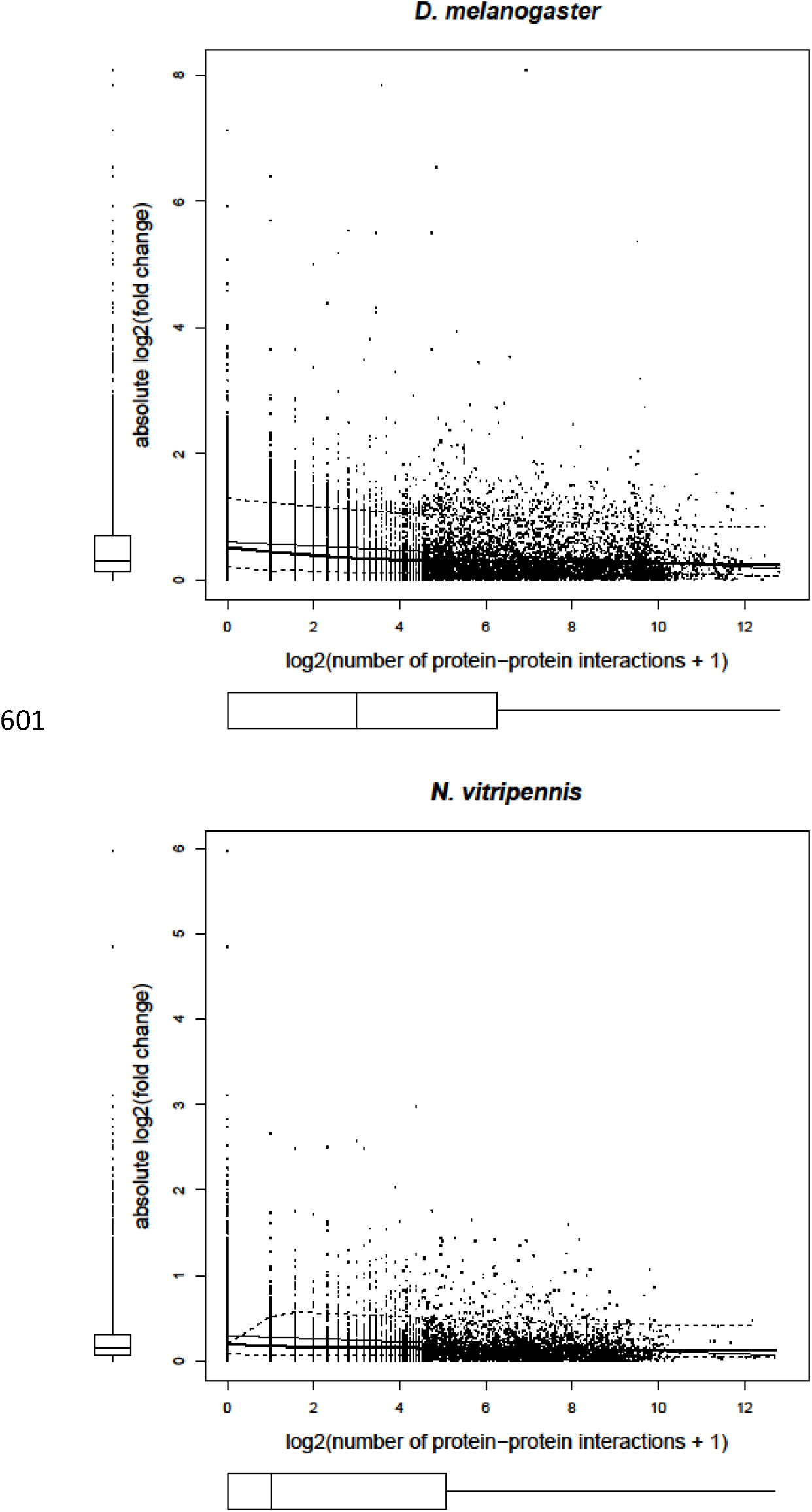
Scatterplots of the log2 fold change of gene expression upon sugar-feeding against the known log2-transformed number of protein-protein interactions for *D. melanogaster* and *N. vitripennis*. There is a significant negative correlation in both species.

## Discussion

### Global gene expression patterns

The set of differentially expressed genes upon sugar feeding in both species were enriched for a broad spectrum of GO-terms, showing that both species adjust their gene regulation within four hours after feeding on a sugar solution. The strong positive correlation of the gene expression levels of conserved (single-copy) genes shows that these patterns are maintained over long evolutionary timeframes (Diptera and Hymenoptera diverged about 340 million years ago (Misof et al. 2014)) and validates our subsequent comparisons of conserved metabolic pathways between *D. melanogaster* and *N. vitripennis*.

### Transcriptional response to sugar-feeding in D. melanogaster

*D. melanogaster* accelerates its fatty acid synthesis after sugar-feeding, as indicated by the upregulation of the key enzyme *fatty acid synthase 1*. This was expected, as *D. melanogaster* is known to start lipogenesis shortly after sugar-feeding (Zinke et al. 2002). *Lipid storage droplet 1*, involved in storage of lipids in the fat body, was upregulated concomitantly, as would be expected when lipid production is increased. The downregulation of genes related to catabolism of amino acids upon feeding indicate that the flies may have used amino acids as fuel in the starvation treatment.

Nine non-coding RNAs were down-regulated upon sugar-feeding. The regulatory role of ncRNAs is a dynamic field of research (Rastetter et al. 2015; Su et al. 2016; Zhao et al. 2016), without generalized predictions of the function of specific ncRNAs. We show here that these nine ncRNAs are either co-regulated with the metabolic genes of *D. melanogaster* or regulating them. Note that our library preparation methods excluded all miRNAs. The transcription factor *sugarbabe* was strongly upregulated in our experiment. In an earlier microarray study this transcription factor was shown to be upregulated in *D. melanogaster* larvae shortly after sugar-feeding (Zinke et al. 2002).

### Transcriptional response to sugar feeding in N. vitripennis

Fatty acid synthesis was not activated in response to sugar feeding in *N*. *vitripennis* as indicated by the lack of a response of the three *fatty acid synthases* in the sugar-fed treatment. However, several other genes that play a direct or indirect role in fatty acid synthesis were upregulated. The gene *acetyl-CoA carboxylase* adds a carboxyl group to acetyl-CoA yielding malonyl-CoA which is required for fatty acid synthesis. High levels of malonyl-CoA also inhibit the activity of *carnitine acyltransferase I*, preventing fatty acid transport to the mitochondrion, thereby limiting β-oxidation. *Glucose-6-phosphate dehydrogenase* produces NADPH^+^, required for anabolic processes like fatty acid synthesis. Activity of *ATP-citrate lyase* is indicative of acetyl-CoA transport from the mitochondrion to the cytoplasm, generally implicated in fatty acid synthesis. The upregulated *citrate transporter* potentially facilitates this transport. Citrate in the cytoplasm stimulates the activity of *acetyl-CoA carboxylase*.

These changes in gene expression in response to sugar-feeding indicate that most of the response of the lipid metabolism is intact, which contrasts with the phenotypic lack of lipogenesis. This paradox could be explained by evolutionary constraints on these enzymes: many enzymes have more than one function (pleiotropy), and the regulatory changes required to decouple all these processes might require many mutations.

A number of other pathways linked to carbohydrate metabolism were differentially regulated upon sugar-feeding. For example, gluconeogenesis was decelerated as indicated by a downregulation of *phosphoenol pyruvate carboxykinase* transcripts. A reduction of gluconeogenesis is expected to be concomitant with a reduction in ketogenesis. However, the upregulation of *hydroxymethylglutaryl-CoA synthase 1* indicates that ketogenesis was accelerated. Catabolism of acetyl-CoA through increased ketogenesis might be linked to loss of lipogenesis in *N. vitripennis*. An earlier study using qPCR to assess gene transcriptional responses to sugar feeding in *N. vitripennis*, found largely congruent results for key genes involved in carbohydrate, fatty acid, and glycerolipid metabolism, including *acetyl-CoA carboxylase, ATP citrate lyase, glucose-6-phosphate dehydrogenase* and *phosphoenolpyruvate carboxykinase* (Visser et al. 2012). Four non-coding RNAs were upregulated and two were downregulated in *N*. *vitripennis*. This suggests that these non-coding RNAs in *N*. *vitripennis* could be involved in regulating its diverging metabolic response to sugar-feeding. Four transcription factors were upregulated and two were downregulated in our experiment. As with the non-coding RNAs, we lack information on their targets and mechanism. It is possible that one of these, or their combined effects, regulate the decoupling of lipogenesis from sugar metabolism.

### Comparison between N. vitripennis and D. melanogaster

It is unclear how the outliers in the comparison of expression levels of single-copy orthologs relate to differences in lipogenic abilities between the studied species. Similarly, it is unknown what the differences in expression patterns of non-coding RNAs and transcription factors mean. Nonetheless, our data indicate that *D. melanogaster* accelerates fatty acid synthesis upon sugar-feeding, while some key components of this pathway in *N. vitripennis* lack a response. In the carbohydrate metabolic pathways, *D. melanogaster* showed many differentially expressed genes in fructose, galactose and sucrose metabolism, while *N*. *vitripennis* had no genes differentially expressed in these pathways. By contrast, differentially expressed genes in *N*. *vitripennis* were involved in the tricarboxylic acid (TCA) cycle, propanoate and butyrate metabolism, and pyruvate metabolism. Moreover, there are contrasting responses in the amino acid metabolisms, lipid metabolic pathways and in the pathways of the metabolism of co-factors and vitamins. Although we currently cannot exclude temporal differences in the patterns of gene expression, these results suggest that these species use a different method of metabolizing dietary sugar and have a divergent response to sugar-feeding overall.

### Differential expression correlates to gene pleiotropy

The negative correlation between fold change and the number of PPI indicates that highly pleiotropic genes are constrained in their extent of up- or downregulation, which is particularly relevant when the insect’s metabolism is under selection. Many genes related to fatty acid metabolism are positioned at the high end of the pleiotropy spectrum, which means that most genes are rather constrained in their change in gene expression. We expected that the regulatory reorganization of fatty acid and acetyl-CoA metabolism of *N. vitripennis* is likely to be directed by genes of low pleiotropy. A good candidate with zero known PPI would be *malonyl-CoA decarboxylase* (LOC100120093). This gene encodes the enzyme that converts one of the substrates for *fatty acid synthase*, malonyl-CoA, to acetyl-CoA. It has a high constitutive expression in *N. vitripennis* in both feeding treatments (logCPM=7.415). The high expression of this gene could therefore potentially deplete available malonyl-CoA, which would impede fatty acid synthesis. *Malonyl-CoA decarboxylase* has been lost in *D. melanogaster*. Other potential candidates having low PPI that likely underlie loss of lipogenesis are enzymes catabolizing acetyl-CoA, or otherwise disposing of it, such as genes involved in ketogenesis and the TCA cycle. Indeed, we show *HMG-CoA synthase 1*, an intermediate step in ketogenesis, to be upregulated in *N. vitripennis* upon sugar-feeding.

### Conclusion

We characterized the gene expression patterns of the non-lipogenic *N. vitripennis* upon sugar-feeding in order to find clues to the molecular mechanism underlying the evolutionary loss of lipogenesis in this species. Animals feeding on sugar generally upregulate their lipogenic pathways (Towle et al. 1997; Kersten 2001), as we observed for the lipogenic *D. melanogaster*. *N. vitripennis* seems to have evolved a regulatory mechanism that decouples sugar ingestion and lipogenesis. Rather than storing dietary sugar in the form of fat, it uses the high expression of *malonyl-CoA decarboxylase* to counteract the activity of *acetyl-CoA carboxylase* and subsequently direct the acetyl-CoA to ketogenesis. The carbohydrates from sugar-feeding are used in somatic maintenance and as a source of energy for physical activity as parasitoid wasps feeding on sugar live longer and loose less fat reserves (Jervis et al. 2008 and references therein). Catabolizing excess carbohydrates via ketone bodies probably helps to avoid adverse effects of a high glucose diet and simultaneously enable retention of fat stores carried over from the larval stage.

Our results raise the question why these lipogenesis-genes are maintained and expressed at detectable levels in *N. vitripennis* despite apparent lack of function. One explanation could be that the genes involved in lipogenesis have undetected subtle forms of gene degradation that impair enzyme function.

Alternatively, genes involved in lipogenesis could be under purifying selection through their pleiotropic effects. The regulatory mechanism hypothesized here could be a way of blocking lipogenesis in adult wasps, while maintaining the genes’ other pleiotropic functions. This could be tested in a knockdown experiment, for example by using RNAi. Future studies on the regulatory network of lipogenesis would give mechanistic insights in these evolutionary constraints.

## Supporting information

### Supplementary tables

S1. Sample quality measurements, number of recovered reads per sample and mapping success

S2. Differentially expressed genes of *D. melanogaster*

S3. Differentially expressed genes of *N. vitripennis*

S4. Enriched GO-terms of the differentially expressed genes of *D. melanogaster*

S5. Enriched GO-terms of the differentially expressed genes of *N. vitripennis*

S6. Non-plastic genes sorted by residual differences in expression level

S7. KEGG-pathways associated with the differentially expressed genes per species

### Supplementary figures

S1. Overview of the active metabolic pathways in abdomens of *D. melanogaster*

S2. Overview of the active metabolic pathways in abdomens of *N. vitripennis*

## Acknowledgements

Lea van de Graaf helped setting up the feeding treatments. The VUmc provided access to their BioAnalyzer and Riet Vooijs helped with measurements. Peter Neleman and Tjalf de Boer advised on bioinformatics analyses. Bart Pannebakker kindly provided the GO-index for *Nasonia vitripennis*. This work was supported by a grant from The Netherlands Organization for Scientific Research [NWO, VICI grant number 865.12.003].

## Data Accessibility

Raw sequence data have been deposited in the NCBI Short Read Archive under the study accession number SRP127311. Derived data (gene expression levels, GO-term enrichment, comparisons of orthologs, KEGG Pathway reaction-based results) are all included as supplementary material to this manuscript.

## Author Contributions

JE conceived the project and performed the feeding experiments. JM performed most molecular wet-lab activities. ML ran the data analysis and interpreted the results. KK and JE advised on data analysis and interpretation. ML wrote the manuscript under supervision of JE and KK.

